# NeoRdRp: A comprehensive dataset for identifying RNA-dependent RNA polymerase of various RNA viruses from metatranscriptomic data

**DOI:** 10.1101/2021.12.31.474423

**Authors:** Shoichi Sakaguchi, Syun-ichi Urayama, Yoshihiro Takaki, Kensuke Hirosuna, Hong Wu, Youichi Suzuki, Takuro Nunoura, Takashi Nakano, So Nakagawa

## Abstract

RNA viruses are distributed throughout various environments, and most RNA viruses have recently been identified by metatranscriptome sequencing. However, due to the high nucleotide diversity of RNA viruses, it is still challenging to identify novel RNA viruses from metatranscriptome data. To overcome this issue, we created a dataset of RNA-dependent RNA polymerase (RdRp) domains that are essential for all RNA viruses belonging to *Orthornavirae*. Genes with RdRp domains from various RNA viruses were clustered based on their amino acid sequence similarity. For each cluster, a multiple sequence alignment was generated, and a hidden Markov model (HMM) profile was created if the number of sequences was greater than three. We further refined the 426 HMM profiles by detecting the RefSeq RNA virus sequences and subsequently combined the hit sequences with the RdRp domains. As a result, a total of 1,182 HMM profiles were generated from 12,502 RdRp domain sequences, and the dataset was named NeoRdRp. Almost all NeoRdRp HMM profiles successfully detected RdRp domains, specifically in the UniProt dataset. Furthermore, we compared the NeoRdRp dataset with two previously reported methods for RNA virus detection using metatranscriptome sequencing data. Our methods successfully identified most of the RNA viruses in the datasets; however, some RNA viruses were not detected, as in the cases of the other two methods. The NeoRdRp can be repeatedly improved by adding new RdRp sequences and is applicable as a system for detecting various RNA viruses from diverse metatranscriptome data.

## Introduction

Currently, many RNA viruses have been identified by deep sequencing of RNA from diverse environmental samples (i.e., metatranscriptome) (Shi et al., 2018; Sakaguchi et al., 2020; Orba et al., 2021). Indeed, various metatranscriptomic analyses have revealed that RNA viruses, similar to human infectious viruses, such as coronaviruses, orthomyxoviruses, and filoviruses, are present even in non-mammalian species, including fishes (Shi et al., 2018). However, identifying RNA virus-derived sequences from metatranscriptome data is challenging because of the low abundance of viral RNA in sequencing reads and the high nucleotide diversity among RNA viruses (Cobbin et al., 2021). Thus, we have developed a method of double-stranded RNA (dsRNA) sequencing named “fragmented and primer ligated dsRNA sequencing” (FLDS) to enrich the RNA virus sequences in the cellular transcriptome (Urayama et al., 2016, 2018; Hirai et al., 2021). Long dsRNAs are rare in eukaryotic cells; therefore, they are mainly derived from the genomes of dsRNA viruses or replicative intermediates of single-strand RNA (ssRNA) viruses (Kumar and Carmichael, 1998). Moreover, FLDS enables us to obtain entire dsRNA sequences, including both termi, and to reconstruct multi-segmented genomes of RNA viruses based on terminal sequence similarity (Urayama et al., 2016). Accordingly, FLDS can enrich potential RNA virus genomes even though their sequences are dissimilar to known RNA viruses. However, it remains difficult to conclude RNA viral genomes based on FLDS data if genes specific to RNA viruses were not found.

Viruses that utilize RNA as a genetic material form a monophyletic group named *Riboviria* based on taxonomy annotation by ICTV (International Committee on Taxonomy of Viruses, https://talk.ictvonline.org/). *Orthornavirae* is a major kingdom of *Riboviria* containing all the dsRNA and ssRNA viruses — but excluding retroviruses — which harbor RNA-dependent RNA polymerase (RdRp). RdRp genes can be used as markers for most RNA viruses; however, their nucleotide and amino acid sequences vary greatly, and some of them have been identified as fusion genes with other protein motifs (Černý et al., 2014; Wolf et al., 2018; Koonin et al., 2020, 2021). Indeed, it is often challenging to identify RNA virus genes, including the RdRp gene, even in the cases of the potentially complete viral genome sequences obtained by FLDS based on a similarity search using BLAST (Basic Local Alignment Search Tool) with major amino acid databases (e.g., nr database or RefSeq database) (Jia and Gong, 2019; Urayama et al., 2020).

In RdRp domain sequences, there are eight conserved amino acid motifs; notably, the sixth motif containing three amino acid residues, GDD, is highly conserved (Koonin, 1991). Based on these RdRp sequence motifs, Wolf and his colleagues inferred the phylogeny of 4,617 RNA viruses (Wolf et al., 2018). However, the accuracy of the multiple alignment of all RdRp amino acid sequences was disputed due to their diversity (Holmes and Duchêne, 2019; Wolf et al., 2019). Therefore, in order to obtain reliable multiple alignments of RdRp sequences, it is necessary to create multiple sequence alignments for each group defined by a shared high sequence similarity.

This study collected RdRp domain sequences from 23,410 RNA viruses and clustered them based on their similarity to build 1,182 hidden Markov model (HMM) profiles. Since an HMM profile is based on a multiple sequence alignment, it can identify distantly related sequences more efficiently than a pairwise-based search (Mistry et al., 2021). Several bioinformatics approaches using HMM have been devised to identify more distantly related viruses, such as VirSorter2 (Guo et al., 2021) and RVDB-prot (Goodacre et al., 2018; Bigot et al., 2019). Note that since those datasets were based on various proteins in RNA and DNA viruses (Goodacre et al., 2018; Bigot et al., 2019), the protein motifs identified by these programs are not necessarily specific to RNA viruses. Furthermore, this study analyzed RdRp sequences of marine RNA viruses obtained by FLDS and revealed that our method successfully detected RdRp domains more efficiently than the two approaches mentioned above (i.e., VirSorter2 and RVDB-prot). The RdRp construction bioinformatics pipeline and constructed RdRp dataset, including annotation, are publicly available online (https://github.com/shoichisakaguchi/NeoRdRp).

## Materials and Methods

### Data sets

A total of 4,620 amino acid sequences containing RdRp domains annotated by Wolf and his colleagues (Wolf et al., 2018) were downloaded from the NCBI (National Center for Biotechnology Information) GenBank database. Additionally, we obtained 18,790 amino acid sequences of 4,239 RNA viruses from the NCBI Virus database (https://www.ncbi.nlm.nih.gov/labs/virus/vssi/) on June 9, 2021. These datasets are referred to as Wolf-RdRps and NCBI-RNA-virus in this study, respectively.

The UniProt Knowledgebase (UniProtKB) containing manually curated protein sequences and functional information concerning 565,928 genes [UniProtKB Reviewed (Swiss-Prot) 2021-05-11] (Boutet et al., 2016) was used to validate and annotate our dataset. The gene ontology (GO) molecular function of RdRp genes was GO:0003968 (RNA-directed 5’-3’ RNA polymerase activity). In the UniProtKB database, 1,027 out of 565,928 amino acid sequences were categorized as GO:0003968; of these, 836 were validated as RdRp-containing genes (Supplementary Table 1).

In addition, potential RNA virus genome sequences obtained from the marine metatranscriptomic analysis by fragmented and primer ligated dsRNA sequencing (FLDS) were used as a benchmark dataset (Urayama et al., 2018). This dataset contains 228 RdRp domains in 1,143 potentially complete RNA virus genomes. For each genome, open reading frames (ORFs) longer than 30 nucleotides (starting with any sense codon) were predicted using ORFfinder version 0.4.3, with the following options: −ml 30 −s 2 (available at https://www.ncbi.nlm.nih.gov/orffinder/).

### Clustering based on amino acid sequence similarity

Amino acid sequences were clustered using the CD-HIT program version 4.7 (Li and Godzik, 2006; Fu et al., 2012) with the following criteria: a similarity threshold of 60% and word size of 4. A multiple sequence alignment was generated using the L-INS-i program in MAFFT version 7.450 (Katoh and Standley, 2013). To select regions with solid statistical evidence of shared homology for each multiple sequence alignment by replacing an amino acid in the uncertainty regions with a gap (-) using Divvier version 1.0, with the default parameters (Ali et al., 2019). In the multiple sequence alignment, if five or more amino acid positions followed by more than 25% of the sequences consisted of gaps, the region was defined as a boundary between two different domains. Then, the multiple sequence alignment was split at the boundary using our in-house script. Multiple alignments of amino acid sequences were visualized using WebLogo version 3.7.8 with the following options: −c chemistry −U probability (Crooks et al., 2004).

### Creation of Hidden Markov Model (HMM) profiles

For each multiple sequence alignment, an HMM profile was built using hmmbuild, part of the HMMER3 package version 3.1b2 (Eddy, 2011) with default parameters. All HMM profiles were then compressed into an HMMER3 searchable database using hmmpress in the HMMER3 package with default parameters.

### HMM and BLAST search

All HMM searches with HMM profiles were carried out using the hmmsearch program from the HMMER3 package with default parameters and three sequence threshold E-values: “-E 1E-10”, “-E 1E-20” and “-E 1E-30”. Two hits in a sequence with gaps of less than 500 amino acid length were merged using BEDTools version 2.30.0 (Quinlan and Hall, 2010) with a “merge −d 500” option (Supplementary Fig. 1). The BLASTp program from the BLAST+ software suite version 2.12.0 (Camacho et al., 2009) was used with “-evalue 1e-10,” “-evalue 1e-20,” and “-evalue 1e-30” options.

### VirSorter2 and RVDB-prot

VirSorter2 (Guo et al., 2021) was used with the option “--include-groups RNA,” targeting the search group set to RNA viruses only. The HMM profiles obtained from Reference Viral Protein DataBase (RVDB-prot) version 22.0 (Bigot et al., 2019) were used with the hmmsearch program.

### Computer resource

A workstation PC (CPU, Intel Core i9-7980XE 2.60 GHz; Memory, 128 GB; Storage 19 TB SSD) with Linux OS (Ubuntu 18.04.6 LTS) was used for benchmarking each bioinformatics program.

## Results

### Constructing RdRp HMM profiles

The schematic procedure for constructing the RdRp HMM profiles is illustrated in Fig. 1. The details of datasets and software used for the analysis are described in Materials and Methods. First, we obtained the amino acid sequences of 4,620 RdRp genes and those with the RdRp domain selected by Wolf and his colleagues (named Wolf-RdRps) (Wolf et al., 2018). Next, we clustered the 4,620 amino acid sequences based on their sequence similarities and then obtained 3,190 clusters. Clusters containing more than three sequences (262 clusters in total) were used for the HMM profile construction. For each cluster, the amino acid sequences were aligned. Since genes including RdRp domain may contain multiple other domains, unreliable regions in the multiple sequence alignments were replaced with gaps. Then, in the multiple sequence alignment, if five or more amino acid positions followed by more than 25% of the sequences consisted of gaps, the region was split there and divided into two multiple alignments. Subsequently, a multiple sequence alignment consisting of more than nine amino acids was defined as a domain. For each domain, an HMM profile was created, and a total of 426 HMM profiles were obtained. The script for the above-mentioned bioinformatics procedure is available at the website: https://github.com/shoichisakaguchi/NeoRdRp.

Furthermore, we conducted comprehensive similarity searches using the 426 HMM profiles with 18,790 amino acid sequences of 4,239 RefSeq RNA virus genomes downloaded from NCBI Virus (named NCBI-RNA-Virus). A total of 7,463 viral amino acid sequences were obtained through the HMM search. The obtained amino acid sequences were combined with Wolf-RdRps, resulting in 12,502 amino acid sequences named “NeoRdRp-seq.” Applying the same procedure for HMM construction (1st round shown in Fig. 1), we clustered the NeoRdRp-seq dataset based on amino acid sequence similarity, resulting in 1,092 clusters (including 8,516 sequences). Note that 2,753 clusters (including 3,986 sequences) containing fewer than three sequences for each were excluded from the HMM profile construction. The 1,092 clusters were processed as the same procedures, and 1,182 domains were extracted. Fig. 2 shows an example of the domain clearly indicating the well-conserved protein motifs, DxxxxD, GxxxTxxxN, and GDD (Koonin, 1991). For each domain, an HMM profile was created, and the 1,182 HMM profiles in total were obtained, which were named “NeoRdRp-HMM” and are available at https://github.com/shoichisakaguchi/NeoRdRp.

**Fig. 1.**
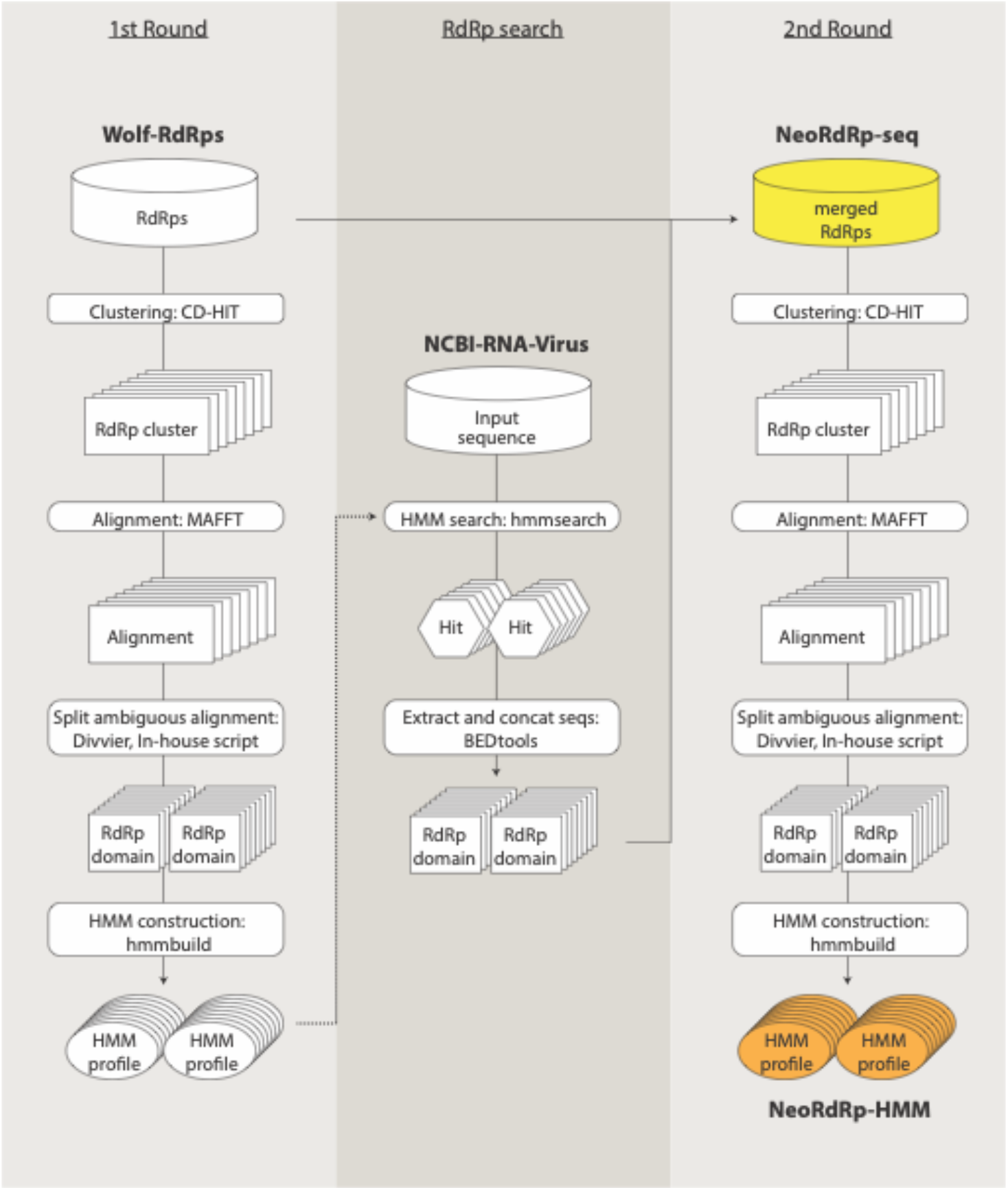
A schematic workflow of NeoRdRp-HMM construction. (1st Round, left) Amino acid sequences containing RdRp domains in the Wolf-RdRps data were clustered based on similarity. For each cluster, a multiple sequence alignment was constructed and processed to create HMM profiles. (RdRp search, middle) The HMM profiles were used to detect the RdRp domain of amino acid sequences in the NCBI-RNA-Virus data. (2nd Round, right) The detected RdRp domain sequences were merged with the Wolf-RdRps data, which were clustered and processed to create HMM profiles for RdRp detection. The dataset names used in this study are in bold. Yellow, NeoRdRp-seq; orange, NeoRdRp-HMM.

**Fig. 2.**
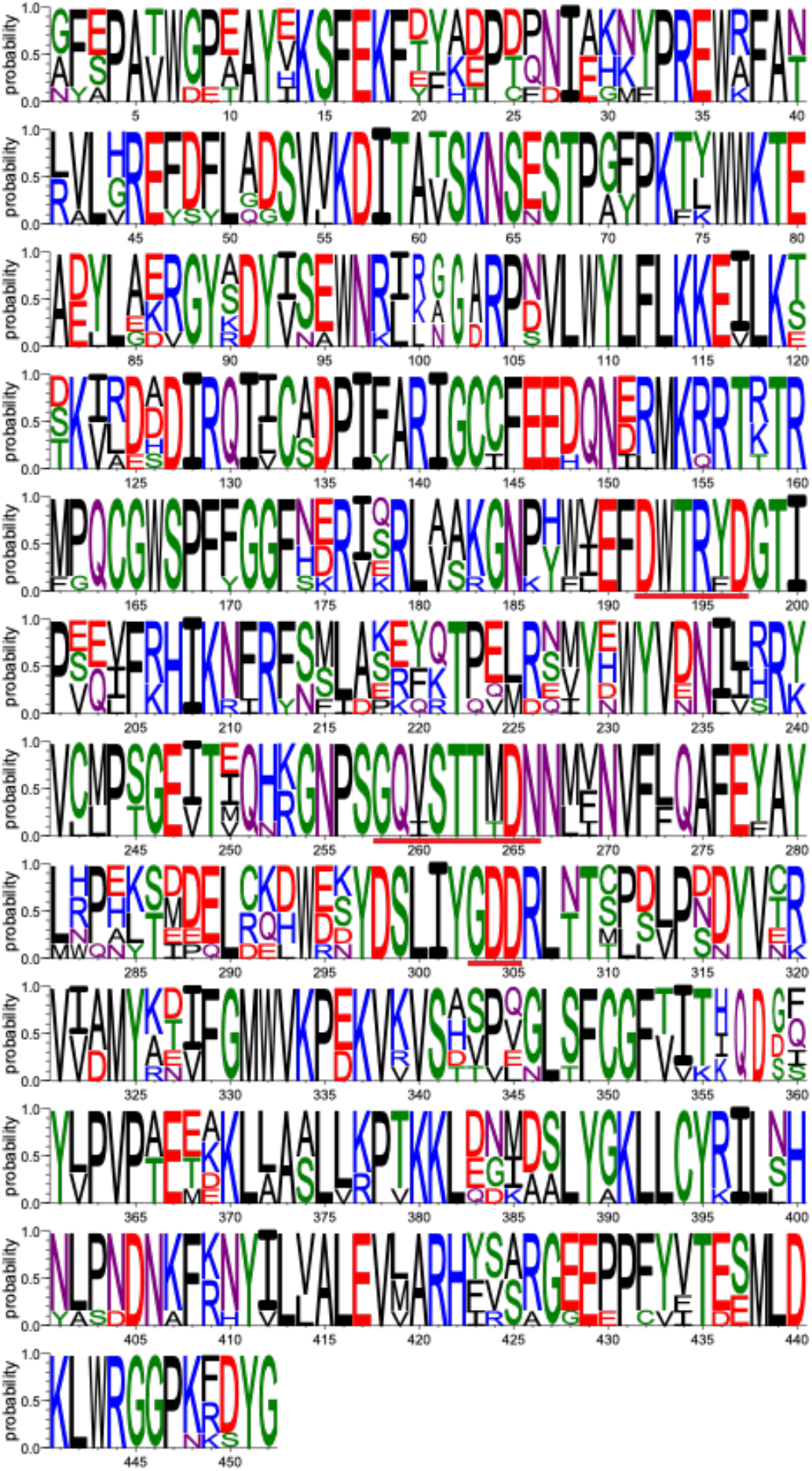
An example of a multiple sequence alignment of amino acid sequences including RdRp domain. The multiple sequence alignment of the HMM profile of 1981.alnta.divvy_RDRP_0.25_923-1375 (including seven sequences) was visualized using Weblogo software. The height represents the probability at each site, and the width is proportional to the number of sequences. Underlines in red indicate the well-conserved DxxxxD, GxxxTxxxN, and GDD motifs in RdRp domain at positions 192–197, 258–266, and 303–305, respectively.

### Evaluating NeoRdRp-HMM and NeoRdRp-seq

To evaluate NeoRdRp-HMM and NeoRdRp-seq, the amino acid sequences and annotation information obtained from the UniProtKB Reviewed database were used. The dataset consisted of 565,928 amino acid sequences, including 836 genes with RdRp domains (see Materials and Methods and Supplementary Table 1). With the 565,928 amino acid sequences as queries, we conducted comprehensive searches using hmmsearch or BLASTp, targeting NeoRdRp-HMM or NeoRdRp-seq, respectively, with three different threshold E-values: 1E-10, 1E-20, and 1E-30. The resulting data are summarized in Supplementary Table 2, and the recall and precision rates calculated from them are summarized in Table 1. For both methods and datasets, the smaller the E-value, the fewer the number of true-positive and false-positive hits there were. In general, NeoRdRp-seq identified more genes with RdRp domain with more false-positive sequences than NeoRdRp-HMM. For example, with the E-value 1E-10 threshold, 813 and 824 out of 836 genes with RdRp domain, and 246 and 1,316 out of 564,418 non-RdRp genes were detected using the NeoRdRp-HMM and NeoRdRp-seq datasets, respectively. Furthermore, our dataset could identify genes with RdRp domain from various RNA viruses, whereas 12 genes with RdRp domain (1.4%) derived from nine Birnaviridae, two Reoviridae, and one Paramyxoviridae were not identified.

**Table 1.**
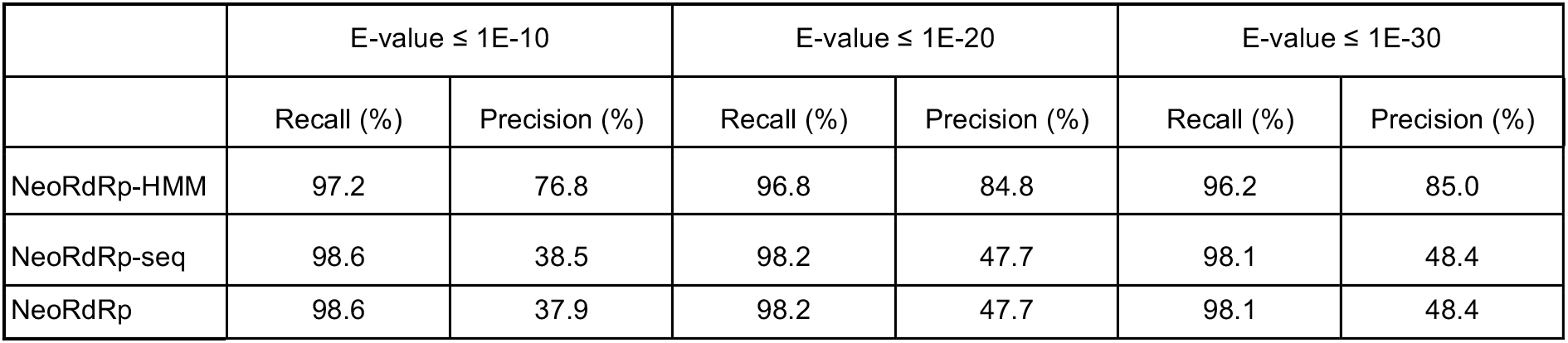
Recall and precision rates of NeoRdRp-HMM or NeoRdRp-seq searches for each E-value threshold.

| | E-value $\leq 1E-10$ | | E-value $\leq 1E-20$ | | E-value $\leq 1E-30$ | |
| --- | --- | --- | --- | --- | --- | --- |
|  | Recall (%) | Precision (%) | Recall (%) | Precision (%) | Recall (%) | Precision (%) |
| NeoRdRp-HMM | 97.2 | 76.8 | 96.8 | 84.8 | 96.2 | 85.0 |
| NeoRdRp-seq | 98.6 | 38.5 | 98.2 | 47.7 | 98.1 | 48.4 |
| NeoRdRp | 98.6 | 37.9 | 98.2 | 47.7 | 98.1 | 48.4 |

We also annotated NeoRdRp-HMM profiles based on the UniProtKB annotation of Gene Ontology (GO; molecular function) with an E-value 1E-10 threshold. Out of 1,182 HMM profiles, 1,012 were evaluated; each hit was scored 1 if “RNA-directed 5’-3’ RNA polymerase activity” (GO:0003968) was included, and 0 if excluded. We then calculated the mean value of the scores for each NeoRdRp-HMM profile, as summarized in Supplementary Table 3. The mean score of the profiles was 0.92, suggesting that most NeoRdRp-HMM profiles annotated using the UniProtKB search were derived from the RdRp domain sequences of RNA viruses. However, we also found that 13 HMM profiles (1.3%) had a score of 0; RNA binding [GO:0003723], ATP binding [GO:0005524], helicase activity [GO:0004386], mRNA methyltransferase activity [GO:0008174], and hydrolase activity [GO:0016787] were shared among the seven HMM profiles. The results indicated that these HMM profiles with a score of 0 detected proteins that interact with RNA but do not contain RdRp domains.

### Comparing RdRp identification

We benchmarked the ability of NeoRdRp-HMM and NeoRdRp-seq to identify RNA viruses using potentially complete RNA virus genome and genome segment sequences obtained by FLDS associated with marine RNA viromes (Urayama et al., 2018). The data comprised 1,143 nucleotide sequences. Among these, 112,108 open reading frames (ORFs) longer than 30 nucleotides (i.e., 10 amino acids) were obtained (see Materials and Methods), including 228 sequences annotated as genes with RdRp domain. The data set of the translated amino acid sequences was named FLDS-data.

Using the FLDS-data, we evaluated NeoRdRp-HMM and NeoRdRp-seq using hmmsearch and BLASTp, respectively. First, NeoRdRp-HMM and NeoRdRp-seq were compared with E-values of 1E-10, 1E-20, and 1E-30; 181, 138, and 123 genes encoding RdRp domain were found by both methods, respectively (Supplementary Fig. 2). Furthermore, with the 1E-10, 1E-20, and 1E-30 E-values, none, 5, and 2 RdRp domain sequences were detected only by NeoRdRp-HMM, whereas 18, 27, and 21 RdRp domain sequences were by NeoRdRp-seq, respectively. The difference in RdRp identification between the two methods could be explained by the advantage of the HMM search in the detection of diversified RdRp domain sequences and the lower coverage of the current HMM models because of the requirement of at least three sequences to create a new reliable HMM profile.

We then merged the results of the NeoRdRp-HMM and NeoRdRp-seq searches into NeoRdRp. Since NeoRdRp-seq may detect large false-positive hits with a low E-value threshold (Table 1), the threshold E-value of the BLASTp search with NeoRdRp-seq was fixed at 1E-30 in this search. VirSorter2 (Guo et al., 2021) and hmmsearch with RVDB-prot (Goodacre et al., 2018) were used for comparison. Note that since VirSorter2 only accepts nucleotide sequences as an input file, the original 1,143 nucleotide sequences of the FLDS-data were used for VirSorter2. First, with a threshold E-value 1E-10 of hmmsearch, we analyzed the 228 genes with RdRp domain using the three methods and detected them as follows: 68 by all three methods, 116 by NeoRdRp and RVDB-prot, one by NeoRdRp and VirSorter2, three by RdRp-prot and VirSorter2, three only by NeoRdRp, 17 only by RVDB-prot, and 20 were not detected by any methods (Fig. 3). Although all hits by NeoRdRp-HMM were also identified by RVDB-prot (Supplementary Fig. 3), viral genomes detected only by NeoRdRp were classified into the family *Flaviviridae, Reoviridae*, and unclassified ssRNA positive-strand viruses based on HMM patterns and genomic structures. However, the viral genomes that the three methods could not detect were recognized as members of *Reoviridae* and *Megabirnaviridae* (Urayama et al., 2018).

**Fig. 3.**
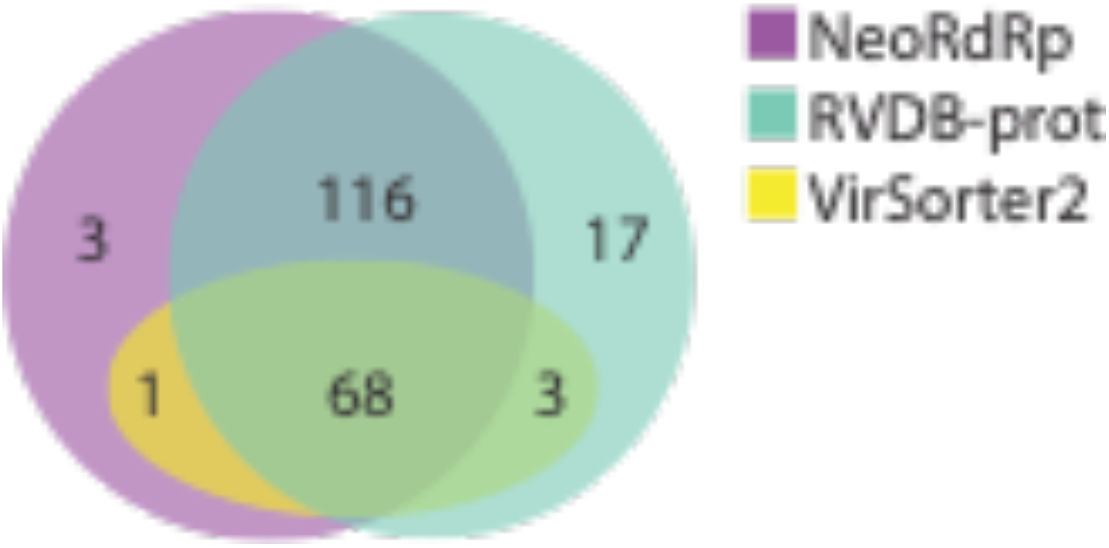
Venn diagram of three different methods for identifying RNA viruses. Annotated marine metatranscriptome data were searched using NeoRdRp (NeoRdRp-HMM and NeoRdRp-seq), RVDB-prot, and VirSorter2. The number in each region indicates the number of hits by the method(s); for example, three were found only in NeoRdRp. Purple, NeoRdRp (NeoRdRp-HMM and NeoRdRp-seq); green, RVDB-prot; yellow, VirSorter2.

We also found 13 possible RdRp-containing sequences in the FLDS-data that were not annotated previously. According to the NeoRdRp-HMM annotation, 13 RdRp domain sequences were similar to those of *Picovirnaviridae*, unclassified viruses, or unclassified dsRNA viruses (Supplementary Table 4). The results suggested that our dataset is useful for detecting RdRps in metatranscriptome data that have been overlooked with previously published bioinformatic pipelines.

We also compared the computational time required to search for RNA viruses in FLDS-data using the three methods (Table 2): hmmsearch using NeoRdRp-HMM and BLASTp using NeoRdRp-seq took 1 min 2 sec and 2 min 56 sec, respectively (i.e., 3 min and 58 sec in total for the NeoRdRp search); hmmsearch using RVDB-prot took 7 min 11 sec; and search using VirSorter took 11 min 45 sec. Furthermore, the database size required for each analysis was 0.37 GB for NeoRdRp, 1.61 GB for RVDB-prot, and 11.3 GB for VirSorter2 (Table 2). The results indicated that the NeoRdRp dataset required a smaller computational capacity than the other methods tested.

**Table 2.**
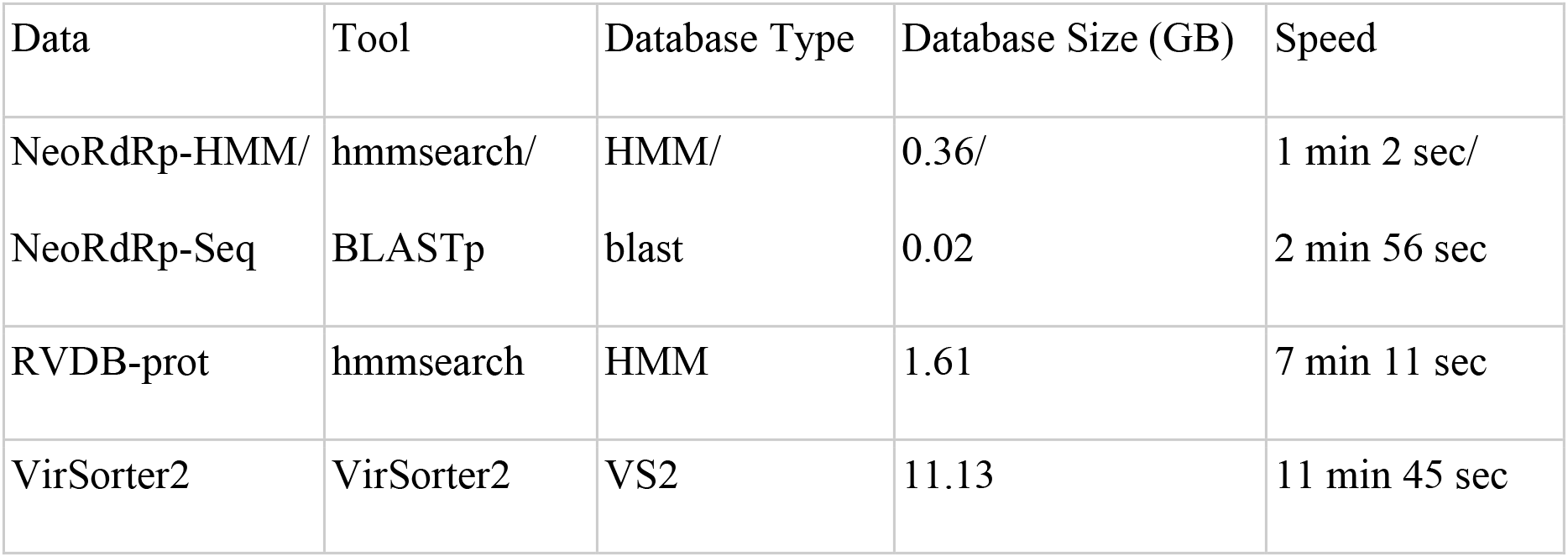
Database size and processing time of three methods for searching for RNA viruses in FLDS data.

| Data | Tool | Database Type | Database Size (GB) | Speed |
| --- | --- | --- | --- | --- |
| NeoRdRp-HMM/ | hmmsearch/ | HMM/ | 0.36/ | 1 min 2 sec/ |
| NeoRdRp-Seq | BLASTp | blast | 0.02 | 2 min 56 sec |
| RVDB-prot | hmmsearch | HMM | 1.61 | 7 min 11 sec |
| VirSorter2 | VirSorter2 | VS2 | 11.13 | 11 min 45 sec |

## Discussion

Nucleic acid contamination from DNA viruses, hosts, and environments is an issue when analyzing RNA viromes. To overcome this issue, techniques to selectively extract and sequence RNA virus-derived nucleic acids, such as the FLDS method (Urayama et al., 2016; Hirai et al., 2021), have been developed. However, it is still challenging to eliminate all contamination from the sequencing data. Thus, in this study, RdRp genes, which are the hallmark of RNA viruses, were comprehensively obtained (NeoRdRp-seq), and HMM profiles of RdRp domains were constructed based on amino acid similarity (NeoRdRp-HMM) (Fig. 1). Using this dataset, we successfully obtained various genes encoding the RdRp domain from the FLDS-data (Fig. 3), while those of some RNA viruses could not be detected (Table 1 and Fig. 3). In addition, although NeoRdRp-seq could be utilized as a complement to RdRp-HMM, a BLASTp search using NeoRdRp-seq generated a relatively large number of false-positive hits (Table 1). This is because an RdRp domain is sometimes encoded in a fusion gene, and NeoRdRp-seq includes non-RdRp domain sequences of such fusion genes. To overcome this problem, repeating updates with additional RdRp sequences is necessary. In principle, as the number of RdRp sequences increases, the number of clusters used for HMM construction also increases, and subsequently, the detection power must increase. We designed the pipeline used for the NeoRdRp-HMM to create enhanced HMM profiles by incorporating newly detected RdRp sequences. The pipeline for HMM construction is available on GitHub (https://github.com/shoichisakaguchi/NeoRdRp). Therefore, users can modify NeoRdRp-HMM using their own sequences. Accordingly, continuing to update NeoRdRp-HMM with successively identified RdRp sequences would reduce undetected RdRp sequences and false-positive hits.

The NeoRdRp dataset was based only on genes containing RdRp domains. Our approach has advantages in the accuracy of RNA virus identification from highly diversified metatranscriptome data, particularly for the presence of known viruses is not expected. Meanwhile, since the NeoRdRp cannot identify non-RdRp genes in RNA viruses, the NeoRdRp dataset can be applied for RNA virus detection and annotation from metatranscriptome data as the following research design; assemble of metatranscriptomic data, subsequent ORF search for each contig, and RdRp domain identification of each ORF using NeoRdRp-HMM. If more RNA virus sequences are required, further searches using NeoRdRp-seq are expected, although it may produce more false-positive hits (Table 1 and Supplementary Table S2) and the process is relatively computationally intensive (Table 2). Based on the combination of these methods and their scores, we can identify contigs containing RdRp domain, suggesting that they are derived from RNA viruses. Furthermore, FLDS can identify multi-segmented genomes of RNA viruses based on terminal sequence similarity (Urayama et al., 2016). Therefore, if one contig is found to contain an RdRp domain, other contigs with a similar sequence on their either termini are strongly implied to be derived from RNA viruses. Further, in these contigs, other viral genes could be detected using other computational programs and datasets, including VirSorter2 and RVDB-prot.

In conclusion, the NeoRdRp can detect various RdRps with a high degree of accuracy (Table 1 and Fig. 3), and the dataset is very compact compared to other methods for RNA virome searches (Table 2). All sequence data, annotations, and bioinformatics pipelines are available on GitHub (https://github.com/shoichisakaguchi/NeoRdRp). Furthermore, by updating the NeoRdRp with new RdRp sequences, it will be possible to find more RdRp sequences with higher accuracy levels from various metatranscriptome data, which may become an essential technique for discovering unknown RNA viruses.

## Acknowledgments

This work was supported by KAKENHI Grant-in-Aid for Scientific Research (C) 20K06775 (to S.S. and S.N.); Early-Career Scientists 20K15685 (to S.S.); Challenging Research (Pioneering) 18H05368 (to S.U., Y.T., and T.Nunoura); and Scientific Research on Innovative Areas 16H06429 (to S.U. and S.N.), 16K21723 (to S.U. and S.N.), 16H06437 (to S.U.), 19H04843 (to S.N.); and by AMED under grant numbers JP19fm0208009 (to S.U.), JP20wm0325004 (to S.U.), and JP21wm0325004 (to S.U. and S.N.). The computing resources were partially supported by the NIG supercomputer at the ROIS National Institute of Genetics, Japan, and SHIROKANE at the Human Genome Center, the Institute of Medical Science, the University of Tokyo, Japan.

---

**Supplementary Figure 1.**
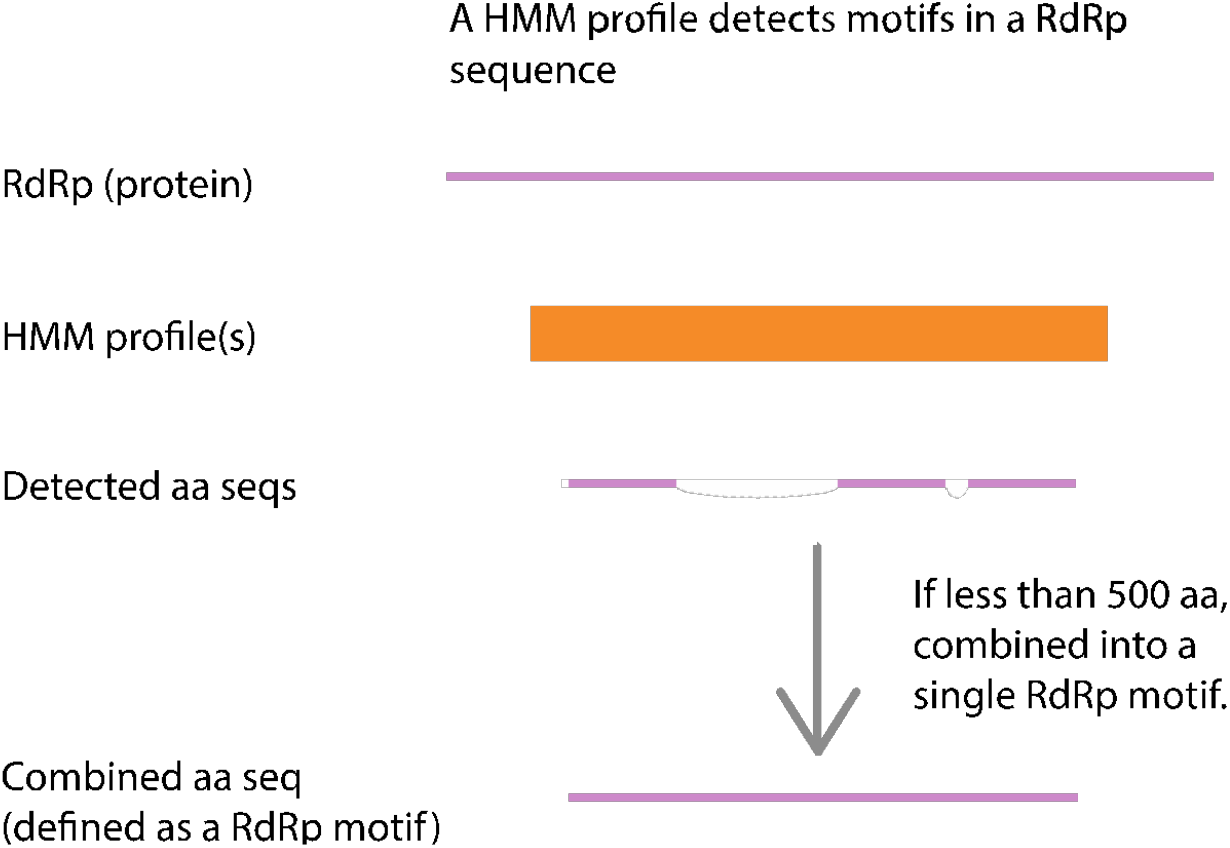
Defining an RdRp domain. When the detected sequence distance was less than 500 amino acid length in the hmmsearch results, we regarded them as a single region as a domain.

**Supplementary Figure 2.**
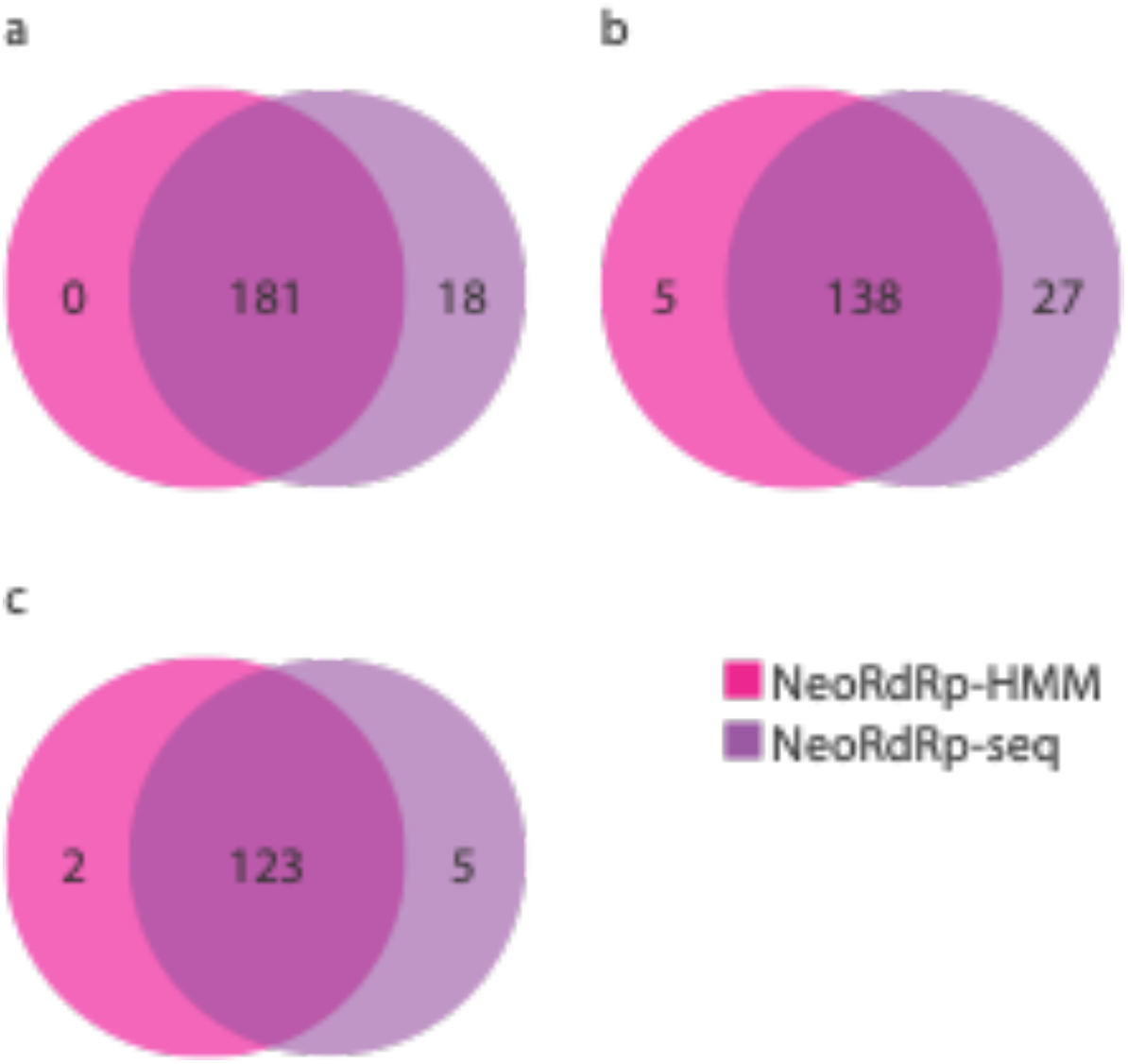
Comparison of NeoRdRp-HMM and NeoRdRp-seq. NeoRdRp-HMM and NeoRdRp-seq with an E-values of (a) 1E-10, (b) 1E-20, and (c) 1E-30 were compared. Pink, NeoRdRp-HMM; purple, NeoRdRp-seq.

**Supplementary Figure 3.**
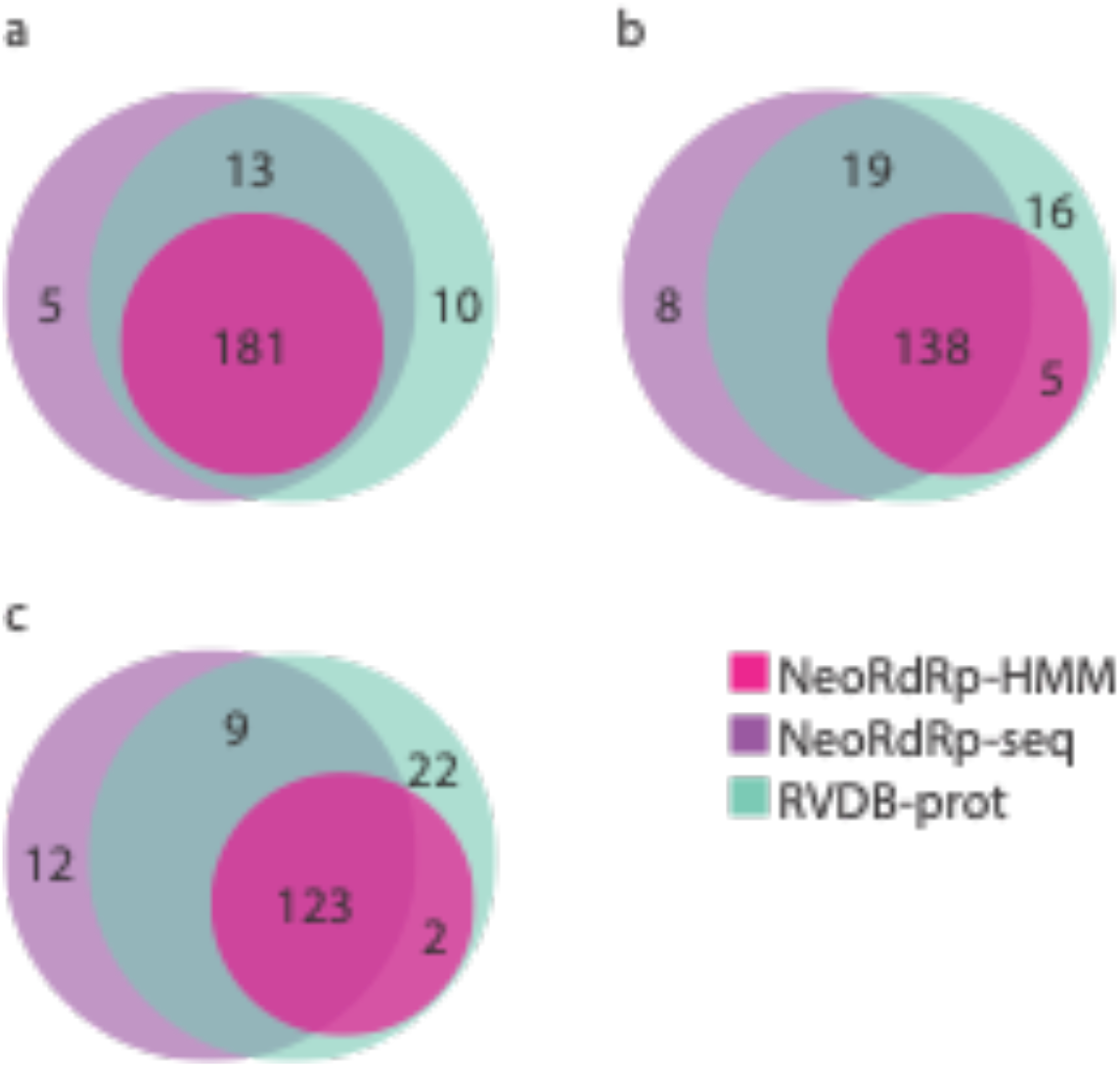
Comparison of NeoRdRp-HMM and RVDB-prot. NeoRdRp-HMM and NeoRdRp-seq with E-values of (a) 1E-10, (b) 1E-20 and (c) 1E-30 were compared. Pink, NeoRdRp-HMM; purple, NeoRdRp-seq; green, RVDB-prot.

**Supplementary Table 1. 836 RdRp genes in the UniProt KB database.**

(provided as an Excel File)

**Supplementary Table 2. NeoRdRp-HMM or NeoRdRp-seq searches for each E-value threshold.**

(provided as an Excel File)

**Supplementary Table 3. Summary of the scores for each NeoRdRp-HMM profile.**

(provided as an Excel File)

**Supplementary Table 4. Unannotated RdRps in metatranscriptome data detected by NeoRdRp.**

